# Recovery from acute SARS-CoV-2 infection and development of anamnestic immune responses in T cell-depleted rhesus macaques

**DOI:** 10.1101/2021.04.02.438262

**Authors:** Kim J. Hasenkrug, Friederike Feldmann, Lara Myers, Mario L. Santiago, Kejun Guo, Bradley S. Barrett, Kaylee L. Mickens, Aaron Carmody, Atsushi Okumura, Deepashri Rao, Madison M. Collins, Ronald J. Messer, Jamie Lovaglio, Carl Shaia, Rebecca Rosenke, Neeltje van Doremalen, Chad Clancy, Greg Saturday, Patrick Hanley, Brian Smith, Kimberly Meade-White, W. Lesley Shupert, David W. Hawman, Heinz Feldmann

## Abstract

Severe COVID-19 has been associated with T cell lymphopenia ^1,2^, but no causal effect of T cell deficiency on disease severity has been established. To investigate the specific role of T cells in recovery from SARS-CoV-2 infections we studied rhesus macaques that were depleted of either CD4^+^, CD8^+^ or both T cell subsets prior to infection. Peak virus loads were similar in all groups, but the resolution of virus in the T cell-depleted animals was slightly delayed compared to controls. The T cell-depleted groups developed virus-neutralizing antibody responses and also class-switched to IgG. When re-infected six weeks later, the T cell-depleted animals showed anamnestic immune responses characterized by rapid induction of high-titer virus-neutralizing antibodies, faster control of virus loads and reduced clinical signs. These results indicate that while T cells play a role in the recovery of rhesus macaques from acute SARS-CoV-2 infections, their depletion does not induce severe disease, and T cells do not account for the natural resistance of rhesus macaques to severe COVID-19. Neither primed CD4^+^ or CD8^+^ T cells appeared critical for immunoglobulin class switching, the development of immunological memory or protection from a second infection.

## Introduction

Several lines of evidence suggest that T cells play important roles in COVID-19^1^. For example, it has been shown that COVID-19 convalescent patients possess both CD4^+^ and CD8^+^ T cells responsive to SARS-CoV-2 antigens^3^. Furthermore, severe COVID-19 is associated with lymphopenia including loss of both CD4^+^ and CD8^+^ T cells^4–6^. However, it is not known whether lymphopenia contributes to severe COVID-19 or is an effect of the disease. Thus, definitive proof of the importance of T cells in recovery from infection and the development of anamnestic responses remains an open question. As an experimental approach to answer this question, we studied adult rhesus macaques that had been depleted of either CD4^+^, CD8^+^ or both T cell subsets prior to infection with SARS-CoV-2 (Fig. 1a). Similar to most adult humans, rhesus macaques become only mildly or moderately affected following infection with SARS-CoV-2, and they do not normally develop acute respiratory distress syndrome^7,8^. Understanding the immunological mechanisms that participate in the resistance of these animals to severe disease is of great interest because it could lead to the rational design of improved vaccines, prophylactics and therapeutics. In this study we focus on the role of T cells in the resolution of acute SARS-CoV-2 infection, and in the development of immunological memory, which provides better protection upon re-infection. It has been shown that rhesus macaques are protected from re-infection^9,10^, but the role of T cells and particularly CD4^+^ T cells in that protection is not yet fully understood^11^.

**Figure 1.**
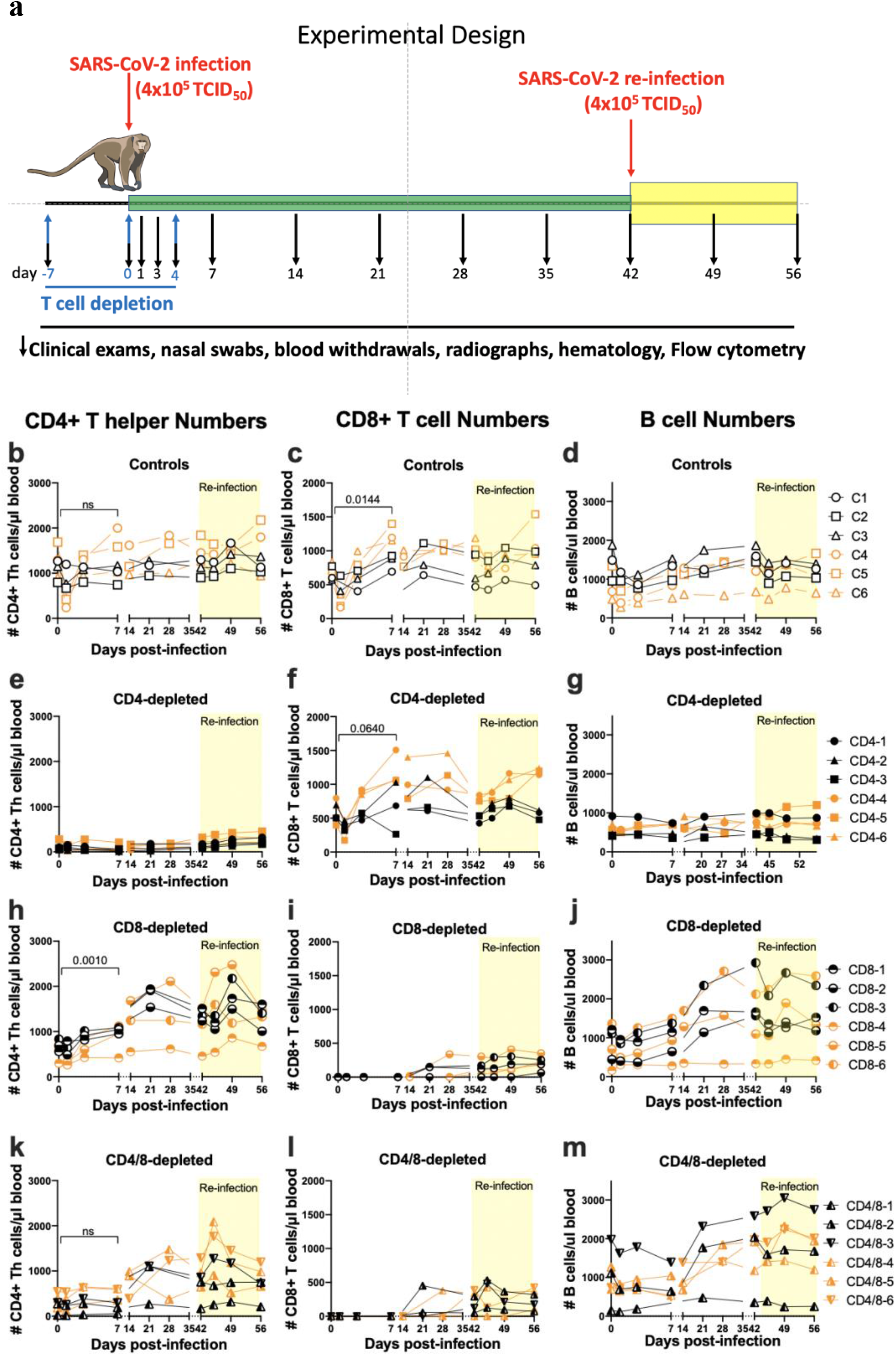
Experimental design and T cell depletions. a. T cell subset-depleting antibodies were administered on days −7, 0 and +4 as indicated by the blue arrows. Infections were done on days 0 and 42 as indicated by red arrows. Blood withdrawals were performed on the days indicated by the black arrows and flow cytometry was used to determine the lymphocyte subset numbers over time. The flow cytometry gating strategies are shown in Supplemental data Fig. 1b. Each symbol represents a single animal throughout. All CD4-depleted animals except CD4-5 were still greater than 90% depleted of CD4^+^ T cells at 7dpi. CD4-5 was 78% depleted. CD4+ Th numbers excluded FoxP3^+^ cells. At 7 days post-re-infection (49 dpi) the animals averaged 81% depleted. All CD8-depleted animals were >99% depleted at 7dpi and remained 78% depleted at 49dpi. The differences between subset numbers at 0 dpi and 7 dpi were calculated by a two-way paired t test. ns = not significant and other p values are shown. Numbers of B cells (d, g, j, m) were determined by flow cytometry using CD45 and CD20 as markers. The numbers of B cells in the CD4-depleted group were significantly lower over time than the controls as determined by mixed effects analysis (p=0.0118).

## Results

All macaques were inoculated with the Washington isolate of SARS-CoV-2 as previously described^7^ and then rested for six weeks. The animals were then challenged a second time as previously. Two separate experiments were carried out, each with three animals per group for a total of six macaques per group. All results from individual animals are labeled with the same symbol throughout: black symbols for animals in the first experiment and orange for those in the second. Findings from the re-infection are highlighted in yellow throughout.

### Lymphocyte responses in normal control animals

Most of the non-depleted control animals showed a rapid but transient lymphopenia with loss of CD4^+^ T helper cells, CD8^+^ T cells and also B cells from the blood, possibly due to homing to lymphoid tissues. CD4^+^ T numbers rebounded to approximately equivalent or higher levels by 7 dpi (Fig. 1b) and CD8^+^ T cell counts were significantly higher at 7 dpi than at day 0 (Fig. 1c) suggesting mobilization or a proliferative response to infection. In support of a proliferative response there was a significant increase in Ki-67 expression at 7 dpi (Supplementary data Fig. 1a). Similar responses were observed following re-infection (yellow shading). B cell numbers in the blood also decreased rapidly after infection and then rebounded over the next several weeks (Fig. 1d).

### Lymphocyte responses in CD4-depleted animals

At day 0 the CD4^+^ T cell depletion in the blood was greater than 90% in all but one animal (#CD4-5 was 78% depleted) (Fig. 1e). No significant increases in numbers were observed in the six subsequent weeks suggesting that little or no immunological priming occurred. Most animals showed slight increases in CD4^+^ T cell counts following re-infection, but it is not known if this was a response to infection or simply reconstitution following depletion (Fig. 1e, yellow shading). The CD8^+^ T cell responses to infection were quite similar to the controls (Fig. 1f) but there was no B cell response apparent in the blood except for a minimal response by animal #CD4-5 following re-infection (Fig. 1f). The reduction of B cell numbers in the CD4-depleted groups compared to the controls over time was statistically significant (p = 0.0118). Thus, depletion of CD4^+^ T cells produced only a minimal impact on CD8^+^ T cell responses but had a significant negative impact on B cell responses.

### Lymphocyte responses in CD8-depleted animals

The expansion of CD4^+^ T helpers in the blood following infection in the CD8-depleted animals was similar or slightly stronger than the controls, although one animal (#CD8-4) failed to respond (Fig. 1h). The CD4^+^ T responses to the second infection also appeared slightly stronger than the controls, possibly a compensatory response to the lack of CD8^+^ T cells (Fig. 1h). The CD8^+^ T cell depletions were >99% effective in all animals and there was no detectable CD8^+^ T cell expansion in the blood for the first two weeks after infection suggesting that minimal or no immunological priming had occurred (Fig. 1i). There was a slight rebound of CD8^+^ T cells in most animals by the day of re-infection but little expansion in the following two weeks. Thus, there was no indication of a significant CD8^+^T cell response to either the first or second infection. All but one animal (CD8-4) had B cell expansions as good or better than the controls (Fig. 1j). The non-responding animal started off the experiment with the lowest CD4^+^ T cell count (Fig. 1h), consistent with the finding that CD4^+^ T cells were important for B cell expansion.

### Lymphocyte responses in CD4/8 dually-depleted animals

The simultaneous depletions of both CD4^+^ and CD8^+^ T cells were less effective for CD4^+^ T cells than the single subset depletions and ranged between 56% and 98% at the time of infection (Fig. 1k). The most strongly depleted animal (#CD4/8-1) displayed little CD4^+^ T cell responsiveness to either the first or second infection. The other five animals showed delayed CD4^+^ T cell responses compared to controls, but increased numbers appeared in the 2 to 4 weeks following infection.

Four of the animals showed rapid CD4^+^ T cell expansion following re-infection indicating that the cells had been primed (Fig. 1k). The CD8 depletions were greater than 99% effective at 7 dpi (Fig. 1l) and both the CD8^+^ T cell responses and B cell responses (Fig. 1m) were very similar to those in the CD8-depleted group. The one animal that failed to make a B cell response was once again the one with the lowest CD4^+^ T cell numbers (#CD4/8-1).

### Virus loads

As previously described for SARS-CoV-2-infected macaques ^7,9^, all animals had high loads of viral RNA in nasal swabs indicative of upper respiratory tract infection. The titers of individual animals in each group are shown in Fig. 2a-d. The RNA loads during the first two weeks of infection were not significantly different between groups, but all control animals cleared infections by 14 days, while the T cell-depleted groups did not totally clear until day 21 or later. The dually depleted animals cleared total viral RNA significantly slower than the controls animals (p = 0.0362, Fig. 2d). In all T cell-depleted groups the second infection was handled better than the first as evidenced by lower viral RNA loads and quicker resolution (Fig. 2a-d, yellow shading). To minimize detection of residual inoculum, nasal swabs were also tested for viral E gene subgenomic mRNA (sgRNA) indicative of replicating virus ^9^. The mean loads of sgRNA (in blue) were lower than the mean loads of total viral RNA but followed similar curves (Fig. 2e-h). Again, all control animals became undetectable for sgRNA at 7 dpi while all T cell-depleted groups still had positive animals. This delay in virus clearance indicated the involvement of CD4^+^ and CD8^+^ T cells in the rapid resolution of acute infection. However, T cells were not essential for eventual clearance. Viral replication in the lower respiratory tract was assayed by analysis of sgRNA in bronchoalveolar lavages (BAL) at 1 dpi and at 1 day post-re-infection (dpri). At 1 dpi, all but one control animal had high viral RNA loads with no significant differences between groups (Fig. 2i-l). Upon re-infection, the lungs showed significantly reduced viral RNA at 1 dpri (d43) compared to 1 dpi, but most animals in all groups had detectable sgRNA in BAL following re-infection. No significant differences between groups were observed at either time point. One or two animals in each group had detectable viral RNA in rectal swabs but no animals had detectable viral RNA in the blood (data not shown). The improved control of virus in the upper and lower respiratory tracts upon re-infection indicated the presence of anamnestic immune responses, and the similarity of all groups in controlling the second infection indicated that such responses were not dependent on an intact T cell repertoire. We next analyzed clinical signs to determine if the reduced viral loads following re-infection were associated with reduced disease.

**Figure 2.**
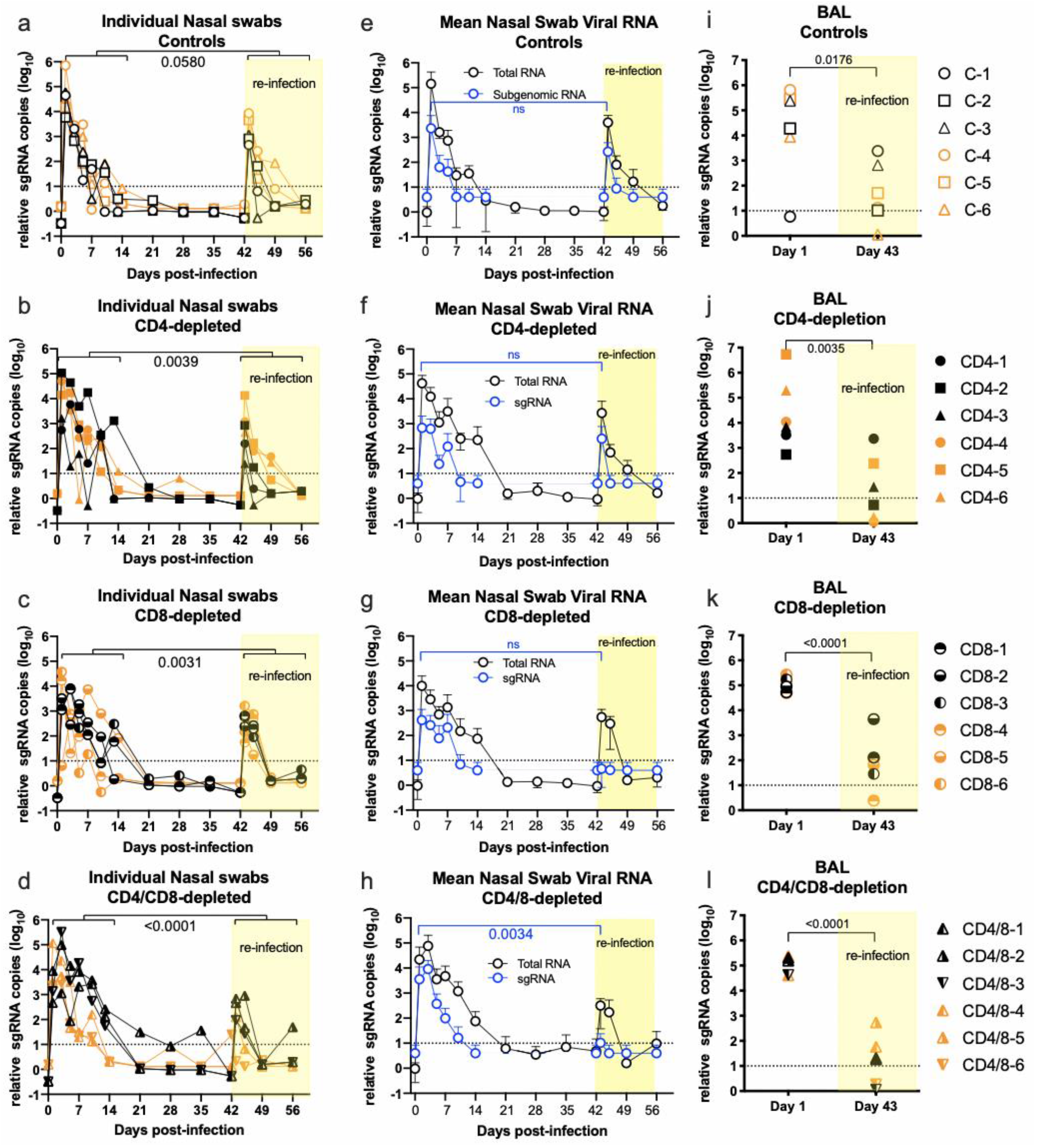
Virus detection from nasal swabs and Broncho-alveolar lavages. (a-d) Each symbol represents the value of viral RNA copies from an individual animal at each time point. The brackets delineate comparisons of cumulative values between the first two weeks after infection with the two weeks after re-infection, and numbers indicate the p values from two-way paired t tests showing significantly reduced virus levels following the second infection (b, c, d) except in the control group (a), which was marginally non-significant. The cumulative RNA titers from the CD4- and CD8-depleted groups for the first two weeks after initial infection were not significantly different than the controls but the CD4/CD8-depleted group had significantly higher titers (p = 0.0362 by one-way Anova with a Dunnett’s post-test). b In (e-h) mean values are shown comparing the total viral RNA data from a-d (black lines) with sub-genomic sgRNA results (blue lines). (i-l) Broncho-alveolar lavage fluids were taken at one day after infection and re-infection. sgRNA was measured from each animal and showed significantly reduced virus replication upon second infection (p values from paired t tests shown in figure).

### Clinical signs

On exam days, each animal was scored in a blinded manner for clinical signs and pulmonary radiographs were performed. The most common clinical signs were reduced appetite, slightly irregular abdominal breathing, ruffled fur and pale appearance (Supplementary data Fig. 3). The cumulative clinical scores for each animal during the first two weeks of infection and re-infection are shown in Fig. 3a-d. All animals developed mild and transient clinical signs during acute infection, and there were no significant differences between the groups. It cannot be excluded that some of the clinical signs were in response to the procedures performed on the animals, which included anesthesia, blood withdrawals, intratracheal virus inoculation and brocho-alveolar lavage. However, upon re-infection the animals underwent the same procedures, and the mean clinical scores in all groups were significantly reduced compared to primary infection (Fig. 3a-d, yellow shading). Pulmonary radiographs showed that all animals developed minor lesions in at least one lobe of the lungs during the first infection (Fig. 3e-h), and the scores were significantly lower after re-infection except in the CD8-depleted group. There were no statistically significant differences in the radiological findings between the control animals and the depleted animals at any time points. We thought it was worth noting that all of the control animals and four of six CD4-depleted animals developed a rapid but transient blood neutrophilia following the first but not the second infection (Fig. 3i). Neutrophilia may result from the stress of handling the animals, but no significant neutrophilia was observed following the first infection in either the CD8-depleted or dually depleted animals (Fig. 3k, l). These findings suggested that neutrophilia was CD8^+^ T cell-dependent. As a whole, these results indicated that neither CD4^+^nor CD8^+^ T cells appeared critical for either recovery from the first infection or improved control of the second infection. Furthermore, there was no evidence that T cell-mediated immunopathology was involved in the disease. The decreased clinical signs following the second infection indicated the induction of an anamnestic immune response that appeared to be largely independent of T cells. Thus, it was of interest to next investigate the antibody responses.

**Figure 3.**
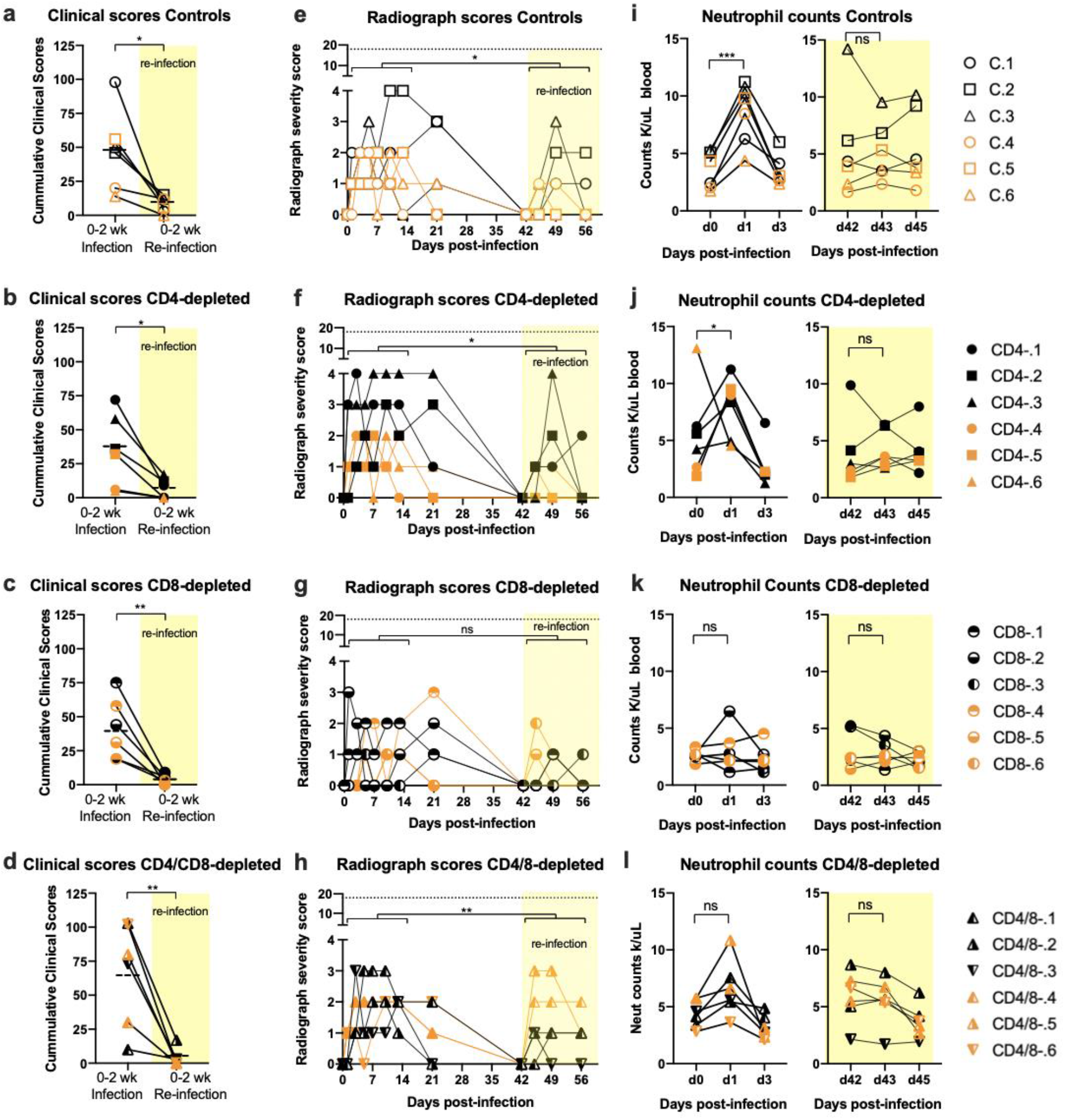
Clinical and radiograph scores and neutrophil counts. (a-d) Clinical signs were scored in a blinded manner using a clinical score sheet. Cumulative scores for the two weeks following the first infection were compared with the two weeks following re-infection (yellow shading) using a two-way paired Student’s t test. The dashed lines indicate means. (e-h) Radiographs were scored in a blinded manner for the presence of pulmonary infiltrates by two board-certified clinical veterinarians. Cumulative scores were analyzed as in a-d. The dotted line indicates the maximum score if all lobes were severely affected. (i - l) Neutrophil counts for each animal were taken on blood withdrawal days and significant differences were noted only between days 0 to 3 as shown. ns = not significant, * = p < 0.05, ** = p < 0.01, *** = p < 0.001.

### Antibody responses. Controls

The development of SARS-CoV-2 receptor binding domain (RBD)-binding antibodies was analyzed in a kinetic manner. The first antibody subtype that develops in response to infections is IgM, and all control animals produced RBD-specific IgM responses peaking at 14 dpi (Fig. 4a). During the antibody maturation process, class switch recombination leads to the development of IgG isotype antibodies in a process usually dependent on CD4^+^ T cells. As expected, IgG responses also developed, lagging behind the IgM responses by one to two weeks and peaking in most animals at 4 wpi (Fig. 4b). The IgG titers had waned slightly by 6 wpi. Upon re-infection, IgG titers rose quickly and dramatically with a mean 37-fold increase in titer within one week (Fig. 4b). All control animals also developed virus-neutralizing antibody (nAb) responses with similar kinetics as the RBD-specific IgG responses (Fig. 1C). These results demonstrated a strong anamnestic antibody response.

**Figure 4.**
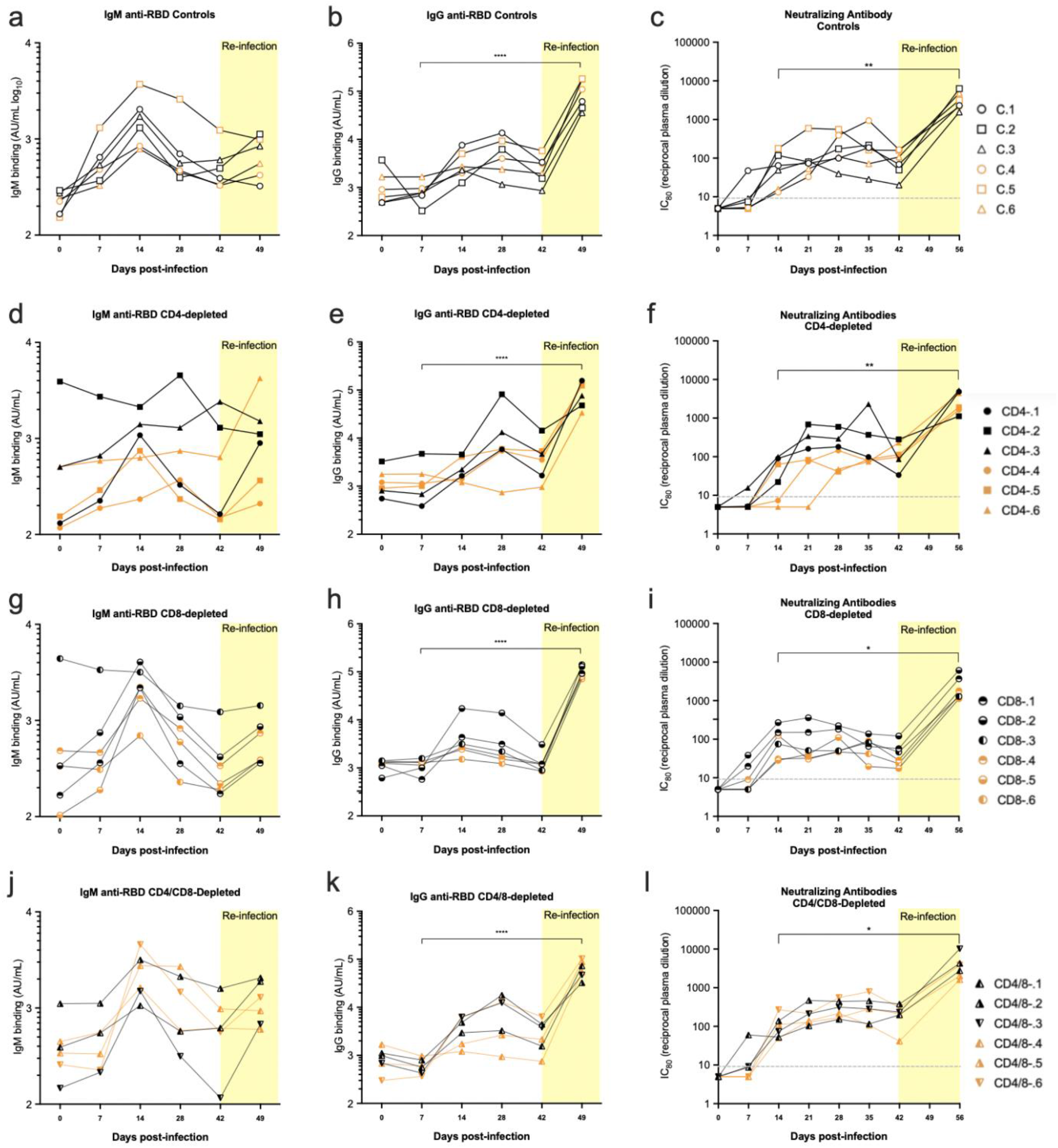
Antibody responses. SARS-CoV-2 spike receptor binding domain-specific IgM (a, d, g, j) and IgG responses (b, e, h, k) were assayed in individual macaques over the course of the experiment using the Mesoscale kit as described in the methods. Each symbol represents a specific animal throughout. Neutralizing antibody titers (c, f, I, l) were measured using a lentiviral pseudovirus expressing the human SARS-CoV-2 spike as described in the method section. Values are reciprocal dilutions that produced an 80% reduction in pseudovirus infection (c, f, i, l). Statistics comparing titers at days 7 dpi and 49 dpi were done using a two-way paired Student’s t test * = p < 0.05, **** = p < 0.0001. Unlabeled comparisons were not statistically significant. Analysis of differences between experimental groups was done by one-way Anova with a Dunnett’s multiple comparisons post-test. The ability of sera from 28dpi to neutralize live SARS-CoV-2 confirmed results from the pseudovirus neutralization assay (Supplemental data Fig. 4).

### CD4-depleted animals

Half of the CD4-depleted animals showed delayed or flat IgM responses (Fig. 4d) and two of them also showed flat IgG responses until re-infection (Fig. 4e). More surprising was that the other four animals developed class-switched IgG responses and all of them developed both virus-neutralizing responses (Fig. 4f) and strong anamnestic responses upon re-infection (Fig. 4e). Of note, one of the animals in the CD4-depleted group (#CD4-2) started the experiment with IgM cross-reactive with RBD (Fig. 4d) as did one of the CD8-depleted animals (#CD8-3). This may have been due to a previous exposure to a different coronavirus. Overall, CD4 depletion delayed and/or dampened IgM and IgG responses in some animals, but all animals developed strong anamnestic IgG responses upon re-infection.

### CD8 and CD4/8-depleted animals

The IgM, IgG and nAb responses of both the CD8-(Fig. 4 g-i) and CD4/8-dually depleted animals (Fig. 4j-l) can be summarized as being very similar to each other and to the controls. No significant effect from CD8-depletions on antibody responses was observed.

In summary, there was no major impact of T cell depletions on the antibody responses even though there was no detectable B cell response in the blood of the CD4-depleted animals (Fig. 1g). The binding and neutralizing antibody titers waned by 42 dpi but rapidly expanded upon re-challenge regardless of T cell depletions. Thus, the memory B cell response may be more critical for protection from re-challenge than standing high titers of antibody.

### Cervical lymph node analysis

T cell levels in blood may not reflect levels in tissues where cells may be more refractory to antibody-mediated depletions. No biopsies were taken during the course of the experiment, but immunohistochemical (IHC) staining was used to examine CD4^+^and CD8^+^ T cells in necropsy tissues at the termination of the experiment (56 dpi). A representative section from a cervical lymph node (LN), which drains the upper respiratory tract is shown for each experimental group (Fig. 5).

**Figure 5.**
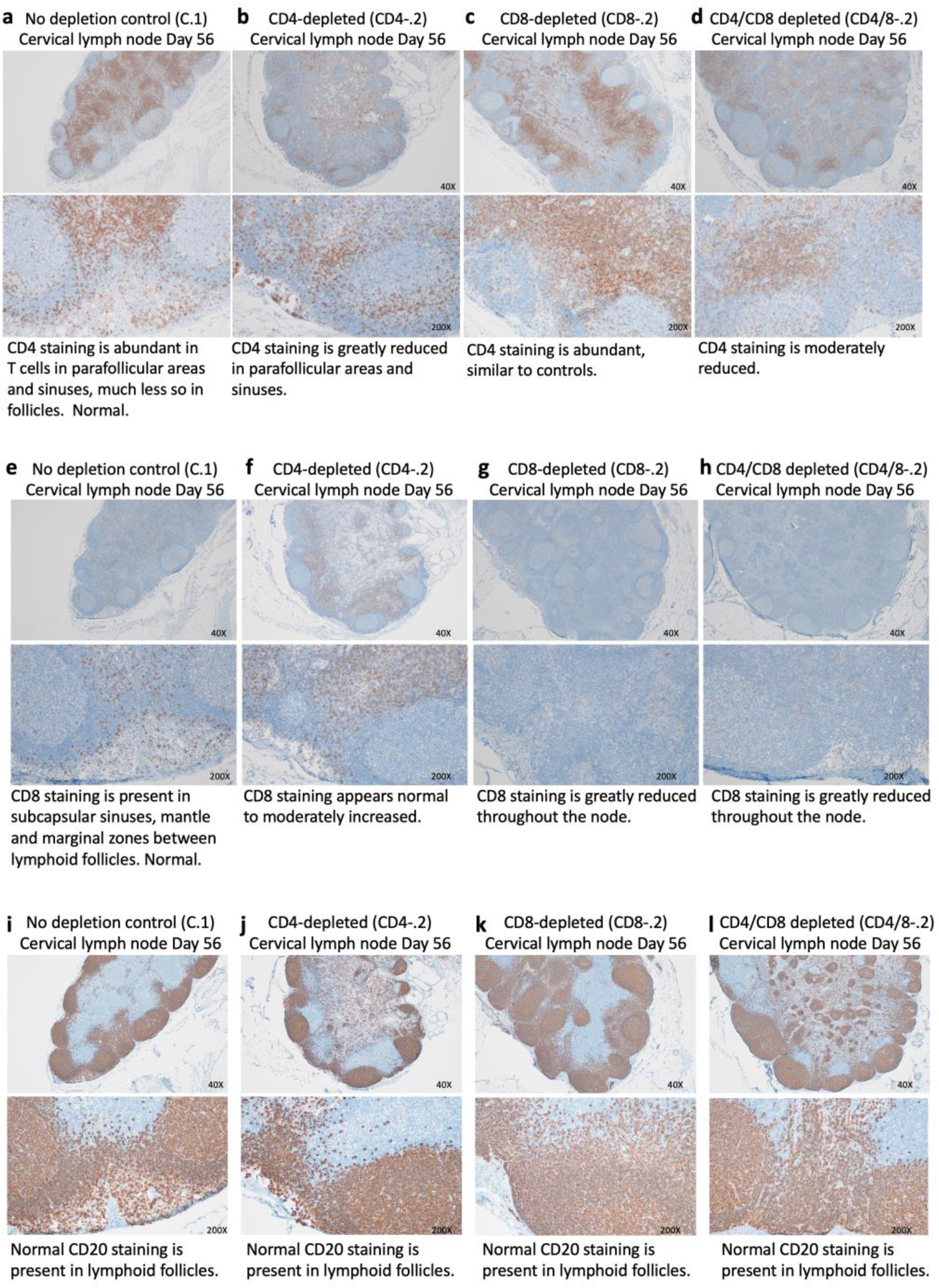
Immunohistochemical staining of cervical lymph nodes for CD4+ and CD8+ T cells and B cells. Representative animals from each experimental group are shown. (a – d) Cervical lymph nodes stained with anti-CD4 antibodies. (e – h) Cervical lymph nodes stained with anti-CD8 antibodies. (i – l) Cervical lymph nodes stained with anti-CD20 antibodies to detect B cells. None of the tissues stained positive for the presence of SARS-CoV-2 at 56 dpi.

### CD4^+^ T cells

Sections of cervical lymph nodes (LN) from controls showed abundant CD4^+^ T cells in periarteriolar lymphoid sheaths and the mantle zone of lymphoid follicles. Representative low and high-magnification sections are shown in Fig. 5a. In comparison, CD4^+^ T cells were greatly reduced in the lymphoid follicles from CD4-depleted animals, although not completely gone (Fig. 5b). CD4^+^ T cell staining appeared normal in the CD8-depleted animals (Fig. 5c), but the CD4/8-depleted animals showed moderately decreased staining (Fig. 5d).

Immunohistochemical staining of spleen sections gave results very similar to the cervical lymph nodes (Supplementary data Fig. 5a, b). These findings cannot distinguish between T cells never depleted and re-seeding of the tissues during the course of the experiment.

### CD8^+^ T cells

Compared to the control (Fig. 5e), the CD8^+^ T cell staining in the CD4-depleted animals appeared normal to slightly increased (Fig. 5f). The LNs from both the CD8-depleted and dually depleted animals showed dramatically reduced CD8^+^ T cell staining (Fig. 5g, h).

### B cells

The LN were also stained with anti-CD20 to detect B cells. Unlike the diminished levels of B cells in the blood of CD4-depleted animals (Fig. 1g), no difference in total CD20 staining was observed in the LN (Fig. 5i-l).

## Discussion

It has been previously reported that rhesus macaques infected with SARS-CoV-2 are resistant to severe disease, similar to most young adult humans ^7^. In this study we investigated the role of CD8^+^ T cells in the natural resistance of macaques to severe COVID-19. The relevance of the results is highlighted by the fact that T cell lymphopenia and T cell dysfunction are associated with severe illness in humans with SARS-CoV-2 infection ^1,2,12,5,13^. Interestingly, we found that depletion of either CD4^+^ and CD8^+^ T cells only slightly prolonged the recovery of macaques from a first infection with SARS-CoV-2 and had little or no impact on recovery from re-infection. This is not to say that T cells do not normally play roles in controlling acute SARS-CoV-2 infections or anamnestic responses, but rather that such roles do not appear critical or that their loss may be compensated by other immune cells in macaques. On the other hand, there was also no evidence that T cells played a pathological role in the development of disease in macaques, as the disease severity in the T cell replete animals was similar to the T cell-depleted animals.

It was unexpected that the CD4-depleted animals would class switch to Ig during primary infection and also mount anamnestic IgG responses upon second infection. The SARS-CoV-2 virion does not contain classic T cell-independent antigens, which are typically rigid arrays of 30 or more repeated epitopes optimally separated by 5-10 nm such as those present on vesicular stomatitis virus ^14,15^. However, there is evidence that SARS-CoV-2 patients mount extrafollicular B cell responses ^16,17^, a type of response that has been shown in mouse studies to produce IgM memory B cells and also IgG class-switched memory B cells in a T cell-independent manner ^18,19^. In that regard most of the mice in all of the groups in this study mounted anamnestic IgM responses (Fig. 4 a, d, g, j), and most also produced inflammatory CXCL-10 (Supplementary Fig. 6), a biomarker associated with severe COVID-19 in humans, and also associated with extrafollicular B cells in COVID-19 patients ^17^. That said, our analysis did not definitively demonstrate the presence of extrafollicular B cells. We cannot exclude the possibility that despite high level depletions in the blood (Fig. 1d), sufficient CD4^+^ T cells remained present in lymphoid tissues to generate immunological help. However, the diminished CD4 staining in cervical lymph nodes at 56 dpi (Fig. 5B) and the strongly ablated B cell responses observed in the blood of the CD4-depleted animals compared to controls (Fig. 1g) argue that the CD4 depletions had a major impact on the T cell repertoire of the animals. B cell activation and stimulation of proliferation typically occur in secondary lymphoid organs where they interact with helper T cells ^20^, but the B cells then disperse and appear in the blood at least transiently, as observed in the control animals but not the CD4-depleted animals (Fig. 1d). It is also possible that help for immunoglobulin class switching was provided by compensatory responses from non-CD4^+^ cells. For example, in the absence of CD4+ T cells, mouse studies with inactivated influenza virus revealed that CD4^−^ CD8^−^ double-negative αβ T cells could provide help for immunoglobulin class switching ^21^. In addition, interferon gamma-producing γδ T cells have also been shown to enable compensatory immunoglobulin class switching ^22^.

In McMahan et al. ^11^ rhesus macaques previously infected with SARS-CoV-2 were depleted of CD8^+^ T cells prior to re-infection, which occurred at 7 weeks after the first infection. In that case all 5 of the depleted animals showed breakthrough virus in nasal swabs whereas only one of the controls did. This finding led to the conclusion that CD8^+^ T cell depletion abrogated the recovery of convalescent macaques. Our study found a slightly prolonged recovery from the first infection but not from re-infection due to CD8^+^ T cell depletion. In our experiments, which employed the same Washington virus isolate and depleting antibody, half of the control animals and half of the CD8-depleted animals showed breakthrough virus in nasal swabs following re-infection as measured by sgRNA (Fig. 2). Experimental differences included the timing of the rechallenge and the CD8 depletions, as well as the addition of the ocular route of challenge in our experiments. While we found less effect on re-infection than the McMahan study, the difference between these two small studies is not statistically different. Both our studies indicate a role for CD8^+^ T cells but a more important role for virus-specific antibodies. It would be of interest in further studies to determine whether T cell immunity in the absence of antibody responses would be sufficient for protection.

Old age in humans plays a significant role in susceptibility to severe disease and death from SARS-CoV-2 infections, and it is well known that advanced age is associated with immunosenescence ^23^ and dysregulated inflammatory responses ^24^ that result in increased vulnerability to infectious diseases. Thus, aged individuals are likely to be considerably less able than adult macaques to compensate for immune deficiencies such as SARS-CoV-2-induced lymphopenia. The oldest macaque in our study was 9 years old, which is not considered to be aged. It has been shown that aged macaques develop lower antibody responses to SARS-CoV-2 infections ^25,26^, so is possible that different results would have been obtained using aged macaques. That said, our results from adult macaques with profound depletion of either CD4^+^ or CD8^+^ T cells leads to the conclusion that neither subset played a critical role in their recovery from acute disease or re-infection. On the other hand, anamnestic antibody responses were strongly associated with recovery from a second infection and reduced disease.

The current studies do not address the requirement for T cells in long-term memory and protection from COVID-19, which will require further experiments. We are currently faced with the emergence of multiple SARS-CoV-2 variants containing mutations in RBD that allow partial or total escape from monoclonal antibodies and reduce vaccine-induced virus neutralization by antibodies ^27, 28^. Furthermore, vaccines based on the original spike sequence appear less protective against the new variants, particularly the South African B.1.351 variant. It is not known whether emerging variants are also evolving to escape T cell responses, but the current results suggest that there may be less evolutionary pressure to escape T cell responses than B cell/antibody responses.

## Acknowledgements

This work was supported by the Intramural Research Program of the National Institute of Allergy and Infectious Diseases of the National Institutes of Health, USA and by Institutional funds from the Division of Infectious Diseases, University of Colorado AMC (MLS)”. Depleting antibodies were obtained from the Non-human primate Reagent Resource. Many thanks to the Rocky Mountain Labs Veterinary Branch animal caretakers for all their hard work. Thanks to Dan Long for his input on the histology and to Emmie de Wit and Vincent Munster for advice and consultation.

## Author Contributions

KJH contributed to experimental concept, design, data analysis and interpretation, figure preparation, statistical analyses and wrote the paper. HF contributed to experimental design, supervision of animal experiments, interpretation of results and manuscript revision. FF contributed to experimental design and scheduling, performed in vivo and in vitro experiments, animal scoring, collection and preparation of experimental samples and performed necropsies. DR performed viral analyses, cytokine analyses, figure preparation and interpretation of results. RJM performed viral analyses. LM designed and performed flow cytometry experiments and analyses and prepared figures. MS, KG, BSB and KM designed and performed neutralizing antibody assays interpreted results and prepared figures. MMC analyzed plasma for virus-specific antibodies and performed viral analyses. AC collected and prepared samples and designed and performed flow cytometry experiments. AO, JL, and KM-W performed animal experiments, collected and prepared samples. CS performed necropsies and histological analyses. RR performed immunohistochemical analyses. PH contributed to scheduling experiments, performed clinical exams, necropsies, bloodwork and radiographs.BS performed clinical exams, necropsies, bloodwork and radiographs. NvD contributed to experimental design. CC performed necropsies. GS performed necropsies. WLS contributed to scheduling and logistics. DWH contributed to experimental design.

## Competing Interests

The authors declare no competing interests.

Materials and Correspondence: Address requests and correspondence to Kim Hasenkrug

## Materials and Methods

### Ethics and biosafety statement

All in vivo experiments were performed in accordance with Animal Study Proposal RML 2020-046-E approved by the Institutional Animal Care and Use Committee of Rocky Mountain Laboratories (National Institutes of Health (NIH)) and carried out by certified staff in an Association for Assessment and Accreditation of Laboratory Animal Care International-accredited facility, according to the institution’s guidelines for animal use, following the guidelines and basic principles in the NIH Guide for the Care and Use of Laboratory Animals, the Animal Welfare Act, and the United States Department of Agriculture and the United States Public Health Service Policy on Humane Care and Use of Laboratory Animals. Rhesus macaques were housed in adjacent individual primate cages allowing social interactions, in a climate-controlled room with a fixed light/dark cycle (12-h light/12-h dark). Commercial monkey chow, treats and fruit were provided twice daily by trained personnel. Water was available ad libitum. Environmental enrichment consisted of a variety of human interaction, manipulanda, commercial toys, videos and music. The Institutional Biosafety Committee (IBC) approved work with infectious SARS-CoV-2 strains under biosafety level 3 conditions. Sample inactivation was performed according to IBC-approved standard operating procedures for removal of specimens from high containment.

### Animals

In the first experiment, 12 adult male rhesus macaques aged 2 1/2 to 5 yrs. were randomly divided into four groups of three. All animals were euthanized and necropsied on day 56 (day 14 post re-exposure). The experiment was repeated with a second group of 12 rhesus macaques including 10 females aged 3 to 10 yrs. old and 2 males aged 4 and 5 yrs. old.

### T cell Depletions

At day −7 (relative to infection) the animals were anesthetized and underwent clinical exams. According to group assignment the macaques received a one-time sq. injection of either 1) (10mg/kg) Rhesus recombinant anti-CD4 depleting antibody (CD4R1), 2) mouse/rhesus CDR-grafted form of the depleting anti-CD8α antibody, M-T807 (each at 10mg/ml prepared in pharmaceutical grade by the NIH Nonhuman Primate Reagent Resource), 3) both CD4R1 and M-T807 or 4) physiological saline. At days −4, 0 and +3 the animals received injections as above except via the i.v. route at 5 mg/kg. as per the provider’s instructions.

### Virus Challenges

On days 0 and 42 the animals were anesthetized and inoculated with SARS-CoV-2 by four routes as previously described ^7^: intratracheal (4 ml), intranasal (0.5 ml each nostril), oral (1 ml) and ocular (0.25 ml each eye). Back titrations for the inocula showed titers of 3.16×10^5^ 50% tissue culture infectious dose (TCID50)/ml for the first challenge of the first experiment, 4.3×10^5^ TCID50/ml for the second challenge, 4.4×10^5^ TCID50/ml for the first challenge of the second experiment and 4.3×10^5^ TCID50/ml for the second challenge of the second experiment. SARS-CoV-2 isolate nCoV-WA1-2020 (MN985325.1)14 (Vero passage 3) was provided by the Centers for Disease Control and Prevention, and propagated as previously described ^7^.

### Clinical Exams and necropsy

Macaques were monitored for clinical signs at least twice daily throughout the experiment using a standardized scoring sheet as previously described ^7^. On exam days (Fig. 1A), clinical parameters such as bodyweight, body temperature and respiration rate were collected, as well as ventrodorsal and lateral chest radiographs. Chest radiographs were interpreted in a blinded manner by a board-certified clinical veterinarian. The total white blood cell count, lymphocyte, neutrophil, platelet, reticulocyte and red blood cell counts, and hemoglobin and hematocrit values were determined from EDTA-treated blood using an IDEXX ProCyte DX Analyzer (IDEXX Laboratories). Necropsies were performed after euthanasia and gross pathology was scored by a board-certified veterinary pathologist. Histopathological analysis of tissue slides was performed by a board-certified veterinary pathologist blinded to the group assignment of the macaques.

### Histology and Immunohistochemistry

Tissues were fixed in 10 % Neutral Buffered Formalin with two changes, for a minimum of 7 days according to IBC-approved SOP. Tissues were processed with a Sakura VIP-6 Tissue Tek, on a 12-hour automated schedule, using a graded series of ethanol, xylene, and PureAffin. Embedded tissues were sectioned at 5 μm and dried overnight at 42°C prior to staining with hematoxylin and eosin. Specific staining was detected using SARS-CoV/SARS-CoV-2 nucleocapsid antibody (Sino Biological cat#40143-MM05) at a 1:1000 dilution, CD4 antibody (abcam cat#ab133616) at a 1:100 dilution, CD8 antibody (Sino Biological cat#10980-T24) at a 1:500 dilution, and CD20 (Thermo Scientific cat#RB-9013) at a 1:250 dilution. The tissues were processed for immunohistochemistry using the Discovery Ultra automated stainer (Ventana Medical Systems) with a ChromoMap DAB kit (Roche Tissue Diagnostics cat#760–159).

### Morphometric analysis

CD4 and CD8 IHC stained sections were scanned with an Aperio ScanScope XT (Aperio Technologies, Inc., Vista, CA) and analyzed using the ImageScope Positive Pixel Count algorithm (version 9.1). The default parameters of the Positive Pixel Count (hue of 0.1 and width of 0.5) detected antigen adequately.

### Thoracic Radiographs

Ventro-dorsal and right/left lateral radiographs were taken on clinical exam days prior to any other procedures. Radiographs were evaluated and scored for the presence of pulmonary infiltrates by two board-certified clinical veterinarians according to a standard scoring system ^29^. Briefly, each lung lobe was scored individually based on the following criteria: 0 = normal examination; 1 = mild interstitial pulmonary infiltrates; 2 = moderate interstitial pulmonary infiltrates, perhaps with partial cardiac border effacement and small areas of pulmonary consolidation (alveolar patterns and air bronchograms); and 3 = pulmonary consolidation as the primary lung pathology, seen as a progression from grade 2 lung pathology. Days 0 and 42 radiographs were taken prior to inoculation, and thus serve as a baseline for each animal. As such, scores for all lung lobes on Day 0 were set to “0 = normal examination.” All subsequent radiographs were compared to the Day 0 radiographs, evaluated for changes from baseline and scored based on the criteria noted above. At study completion, thoracic radiograph findings are reported as a single radiograph score for each animal on each exam day. To obtain this score, the scores assigned to each of the six lung lobes are added together and recorded as the radiograph score for each animal on each exam day. Scores therefore range from 0 to 18 for each animal on each exam day.

### Spike and RBD-binding IgM and IgG

IgM and IgG titers were quantified from serum collected for all animals on Days 0, 7, 14, 28, 42, and 49. Meso Scale Discovery V-PLEX SARS-CoV-2 Panel 1 Kit (Rockville, MD) was used for determining antibody binding for IgG (K15359U) and IgM (K15360U) specific for SARS-CoV-2 S1 RBD. Protocols were followed as per manufacture’s recommendations. All diluted samples fell within standard curve ranges. A Meso Scale Discovery® MESO QuickPlex SQ 120 instrument was used for measuring chemiluminescence with Methodical Minds ™ acquisition software. Analysis was performed on the DISCOVERY WORKBENCH® software (version 4.0). Concentrations are relative to internal controls and were reported as arbitrary units/mL (AU/mL). Graphpad Prism 8 was used for preparation of graphs and statistical analysis.

### Virus-neutralizing antibody assay

Neutralizing antibody titers were determined in plasma samples using a modified lentivirus-based pseudovirion assay ^30^. Plasmids encoding an Env-defective HIV-1 backbone tagged with the nanoluciferase gene (HIV-1_NL_Δ Env-NanoLuc) and an expression construct for the SARS-CoV-2 spike lacking 19 amino acids of the cytoplasmic tail encoding the ER retention motif (CMV-SARS-CoV-2 SΔ19) were kind gifts from Dr. Paul Bieniasz (Rockefeller University). Pseudovirions were prepared in HEK293T cells by co-transfecting 60 μg of HIV-1_NL_ΔEnv-NanoLuc and CMV-SARS-CoV-2 SΔ19 at a 3:2 ratio in T-175 flasks using the calcium phosphate method, then were concentrated by ultracentrifugation in a sucrose cushion ^31^. We utilized A549-ACE2 cells ^32^ as target cells as these cells attached better to the culture plate compared to 293T-ACE2 cells ^30^, allowing for extensive washes given that the nanoluciferase reporter yielded high backgrounds. A549-ACE2 cells were cultured in complete media containing F-12 Ham’s Media (Corning), 10% fetal bovine serum (Atlanta Biologicals) and 1% penicillin/streptomycin/glutamine (Corning). For the Nab assay, a previously-determined pseudovirus titer (300,000 to 400,000 relative light units or RLU per well) were co-incubated with serial 5-fold dilutions of plasma (1:5 to 1:15,625) in 100 μl complete media at 37°C for 1 h in 96-well round-bottom plates. In duplicate, 40 μl of the virus-plasma mixture was combined with 160 μl of complete media containing 10,000 A549-ACE2 cells. The virus-plasma-cell mixtures were plated in white polystyrene plates (Millipore-Sigma) and cultured at 37°C 5% CO2. After 48 h, the spent media in each well were removed, and the cells were washed four times with 200 μl PBS. After the last wash, 100 μl of PBS was dispensed into each well, and mixed with 100 μl of Nano-Glo luciferase substrate working solution (Promega). RLU values were measured in a VictorX5 luminometer (Perkin Elmer). To compute 80% inhibitory concentrations (IC80), the mean RLUs of virus-only wells (n=6) in each plate were set as 100% infection. Duplicate RLUs for each plasma dilution were averaged and normalized against virus-only wells. Best-fit nonlinear regression curves were constructed based on a two-phase decay equation (GraphPad Prism 8), and 80% inhibition values (20% infection) were interpolated. Nab titers were reported as reciprocal plasma dilutions.

Nab titers against live virus were also determined for a subset of plasma samples (28 dpi) in A549-ACE2 cells in a 48-well plate format ^32^. Briefly, 1:20, 1:100 and 1:500 dilutions of plasma were co-incubated at 37°C for 1 h with a nonsaturating dose of the SARS-CoV-2 USA-WA1/2020 strain that would yield ~100,000 copies in the quantitative PCR assay. After 24 h, virus copy numbers were evaluated from culture supernatant using nucleocapsid-specific primers and probes as we previously described ^32^.

### Virus detection

RNA was extracted from swabs and bronchoalveolar lavage using the QiaAmp Viral RNA kit (Qiagen) according to the manufacturer’s instructions and as described ^7^. 5 μl RNA was used in a one-step real-time RT–PCR E assay using the Rotor-Gene probe kit (Qiagen) according to instructions of the manufacturer. In each run, standard dilutions of counted RNA standards were run in parallel, to calculate copy numbers in the samples. For detection of SARS-CoV-2 mRNA, primers targeting open reading frame 7 (ORF7) were designed as follows: forward primer 5’-TCCCAGGTAACAAACCAACC-3’, reverse primer 5’-GCTCACAAGTAGCGAGTGTTAT-3’, and probe FAM-ZEN-CTTGTAGATCTGTTCTCTAAACGAAC-IBFQ. Five μl RNA was used in a one-step real-time RT-PCR using the Rotor-Gene probe kit (Qiagen) according to instructions of the manufacturer. In each run, standard dilutions of counted RNA standards were run in parallel, to calculate copy numbers in the samples. Detection of viral E gene subgenomic mRNA (sgRNA) was performed using RT–PCR as described ^9^. The forward primer was 5’-3’ CGATCTCTTGTAGATCTGTTCTC, the reverse primer was ATATTGCAGCAGTACGCACACA and the probe was FAM-ACACTAGCCATCCTTACTGCGCTTCG-ZEN-IBHQ.

### Flow Cytometry

Immune cell marker analysis was performed on freshly isolated PBMCs following enrichment after a standard Histopaque 1077 (Sigma) centrifugation procedure in 15-mL Leucosep Tubes (Greiner Bio-One) from EDTA blood samples. Briefly, 3 mL of room temperature Histopaque 1077 was added to Leucosep tubes and centrifuged at 1000xg for one minute. Blood was diluted 1:2 in 1X DPBS then transferred into the Leucosep tube and centrifuged at 1000xg for 10 min, RT without the brake. The enriched live PBMC fraction was collected and washed in 1X DPBS and then transferred into plates for staining. Cells were incubated for 30 min with the following cell surface antibodies: BUV661-anti-CD45 (D058-128, BD Biosciences 741657 Lot 0198214), PE-Cy7-anti-CD20 (2H7, BD Biosciences 560735 Lot 0170481), BV786-anti-CD3 (SP34-2, BD Biosciences 563918 Lot 0133645), FITC-anti-CD8 (DK25, Millipore FCMAB176F Lot 3398058), and Pacific Blue-anti-CD4 (OKT4, BioLegend 317424 Lot B258189) in 2% FBS PBS supplemented with Brilliant Stain Buffer (BD Biosciences). Intracellular staining was performed using the Foxp3/Transcription Factor Staining Buffer Set (Thermo Fisher) following the company’s recommendation. Cells were incubated for 30 min with the following antibodies: PE-anti-Foxp3 (259D, BioLegend 3202080 Lot B293618) and AF700-anti-Ki-67 (B56, BD Biosciences 561277 Lot 9315354). Live lymphocytes were gated using a SSC-A and FSC-A gate, by FSC-H and FSC-A to exclude doublets and then by time to exclude artifacts caused by erratic sample flow. Gating strategies for B cells and CD8^+^and CD4^+^ T cells are shown in Supplementary data Figure 4. The multi-parameter data were collected within the BSL-4 using a Cytoflex LX (Beckman Coulter) and analyzed using FlowJo software (version 10.7.1; TreeStar, Inc).

### Statistical Analyses

All statistics were analyzed using Prism Mac OS version 8.4.3 software. The specific tests used for statistical analyses are listed in the figure legends. The p value cutoff for statistical significance was set to 0.05. P values are shown in the figures or figure legends.

---

**Supplementary data Fig. 1a.**
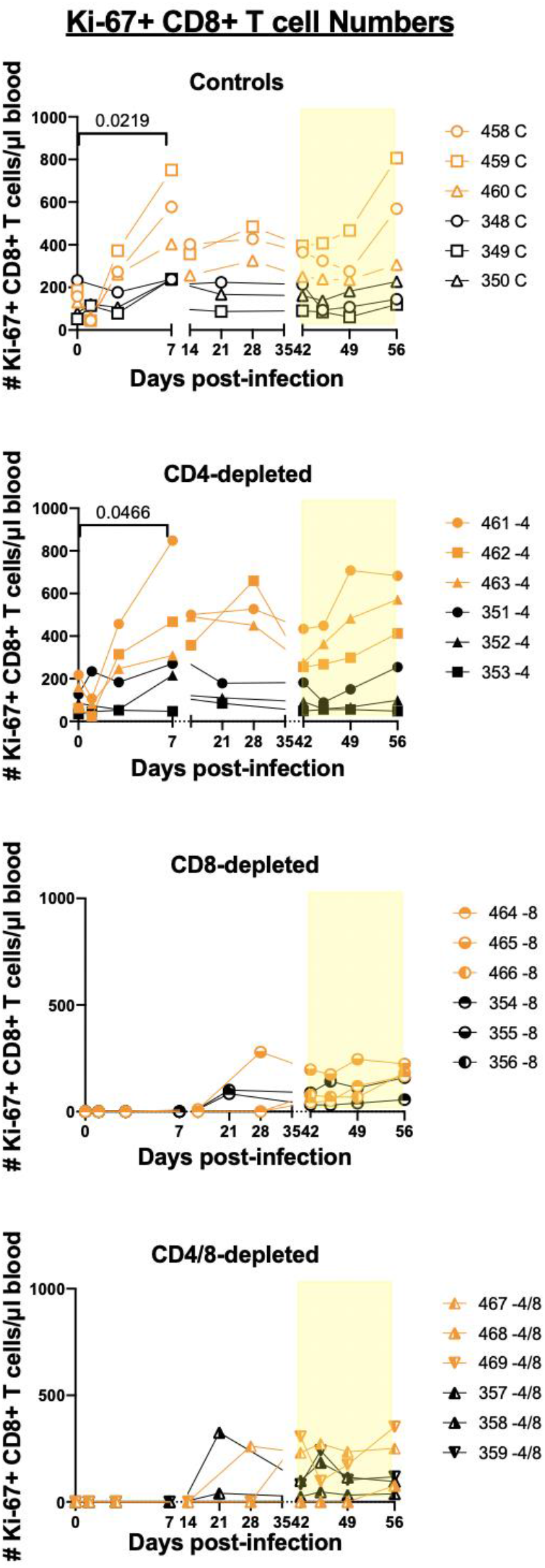
Ki-67 staining of CD8+ T cells

**Supplementary data Fig. 1b.**
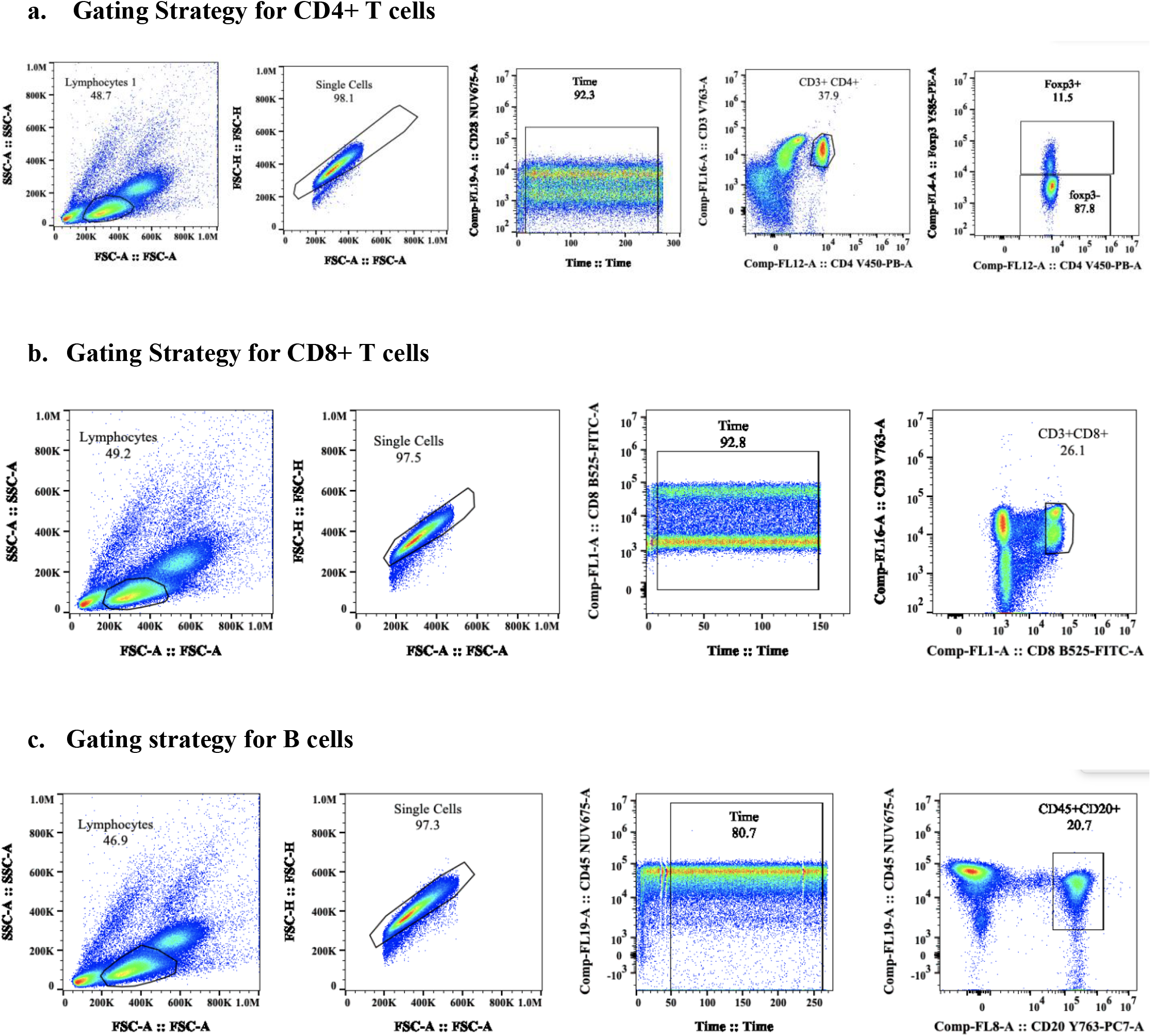
Gating strategies for Flow Cytometry.

**Supplementary data Fig. 3.**
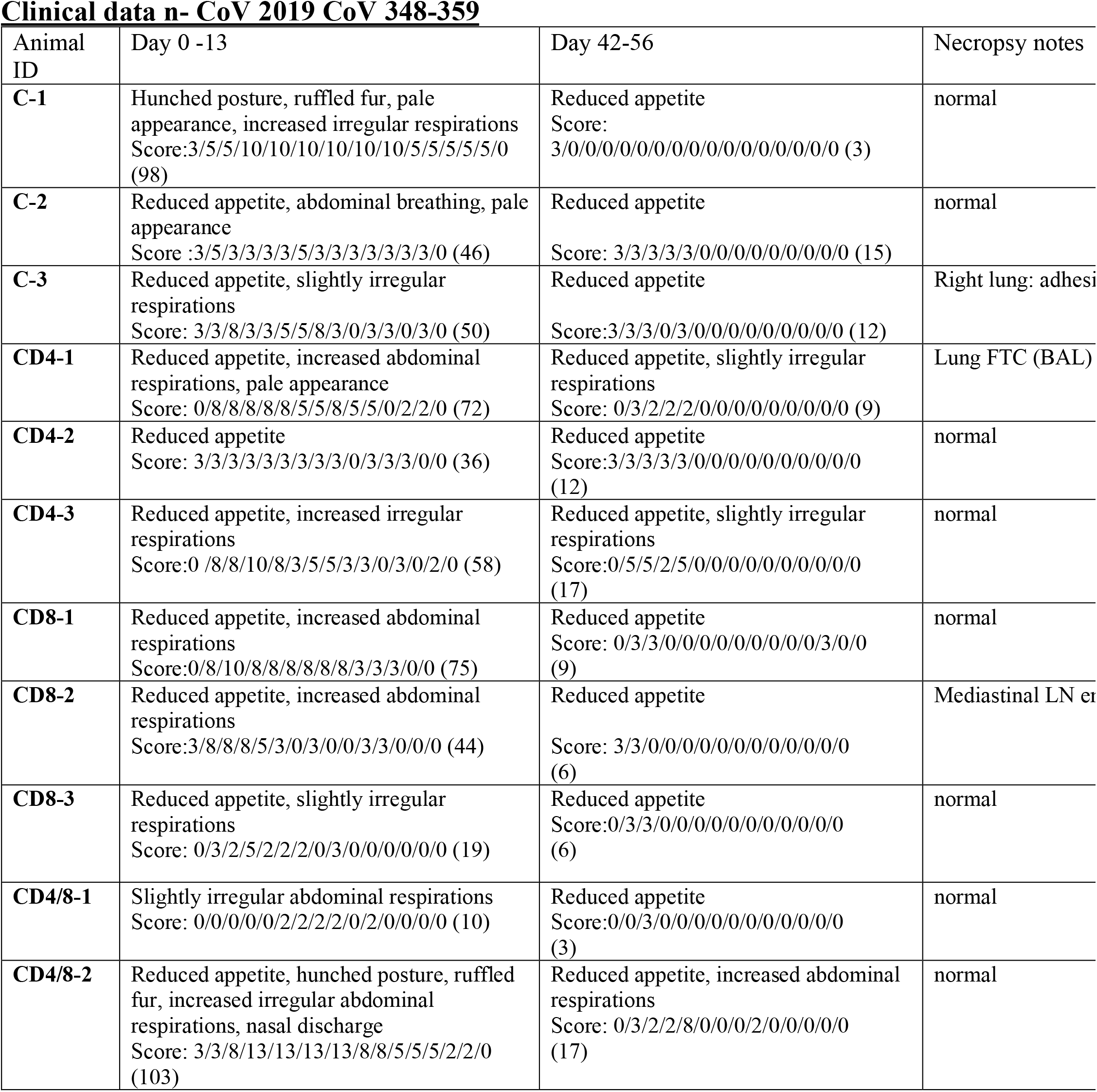
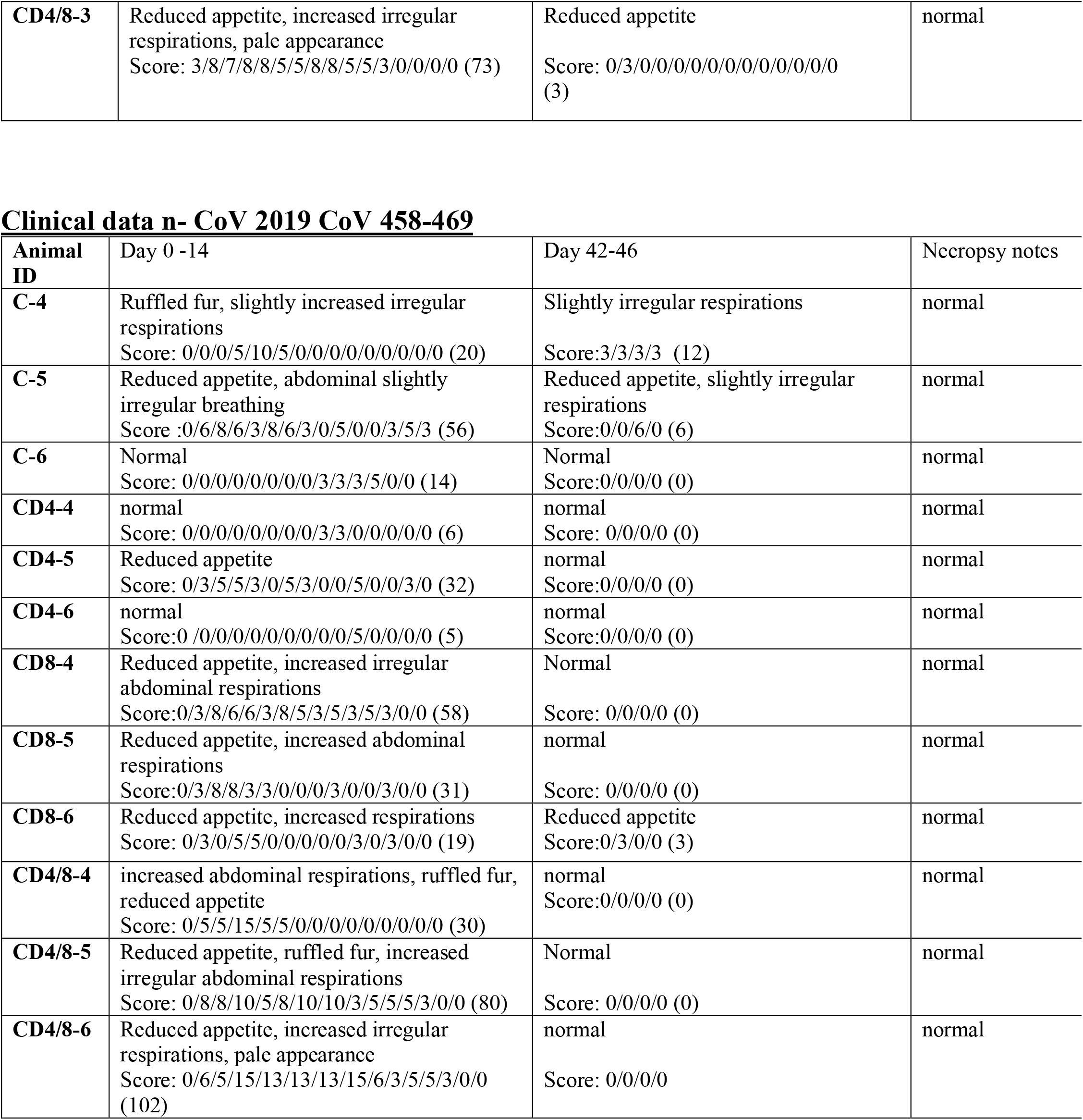

**Supplementary data Fig. 4.**
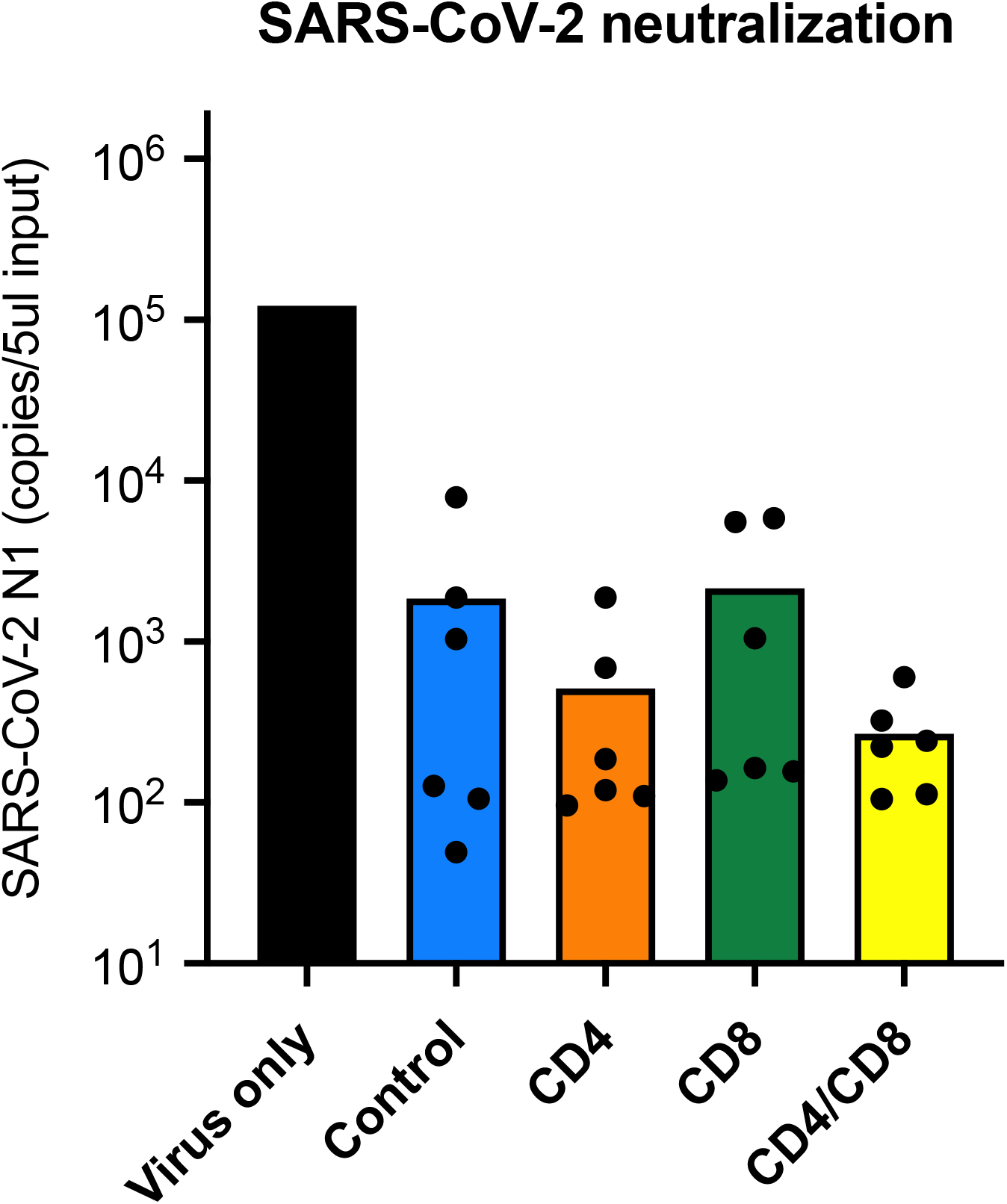
SARS-CoV-2 neutralization assay with live coronoavirus. Day 28 post-infection sera from all NHP were diluted 1:20 and tested for neutralization of SARS-CoV-2. Each dot represents the result from an individual macaque.

**Supplementary data Fig. 5a.**
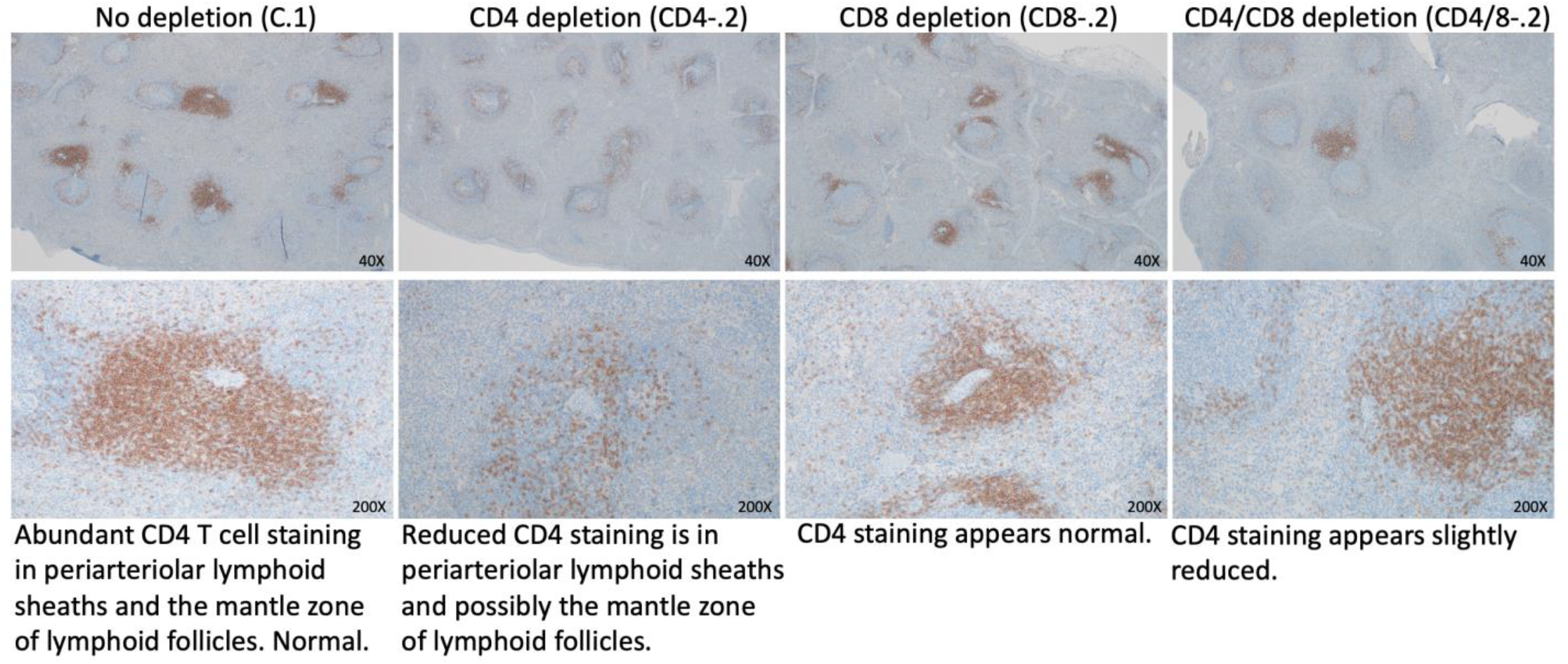
CD4+ T cell staining in spleens.

**Supplementary data Fig. 5b.**
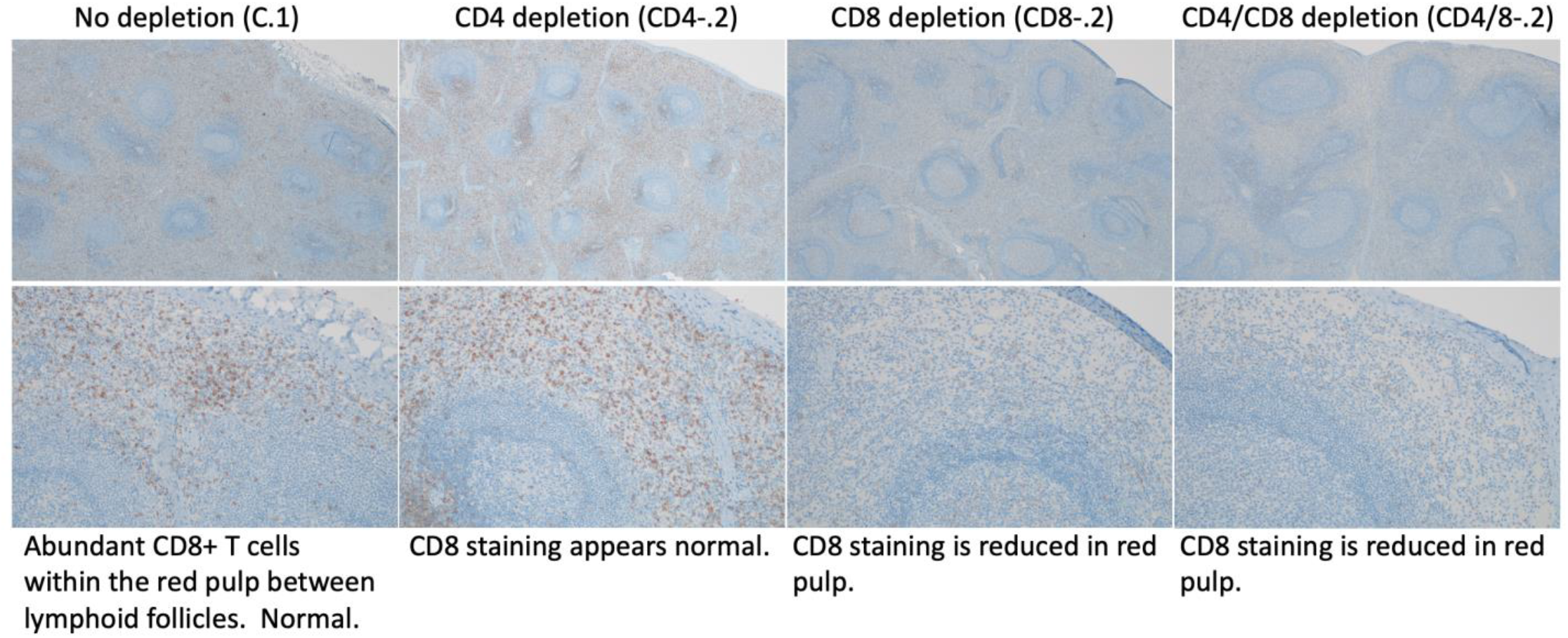
CD8+ T cells in spleen

**Supplemetary Figure 6.**
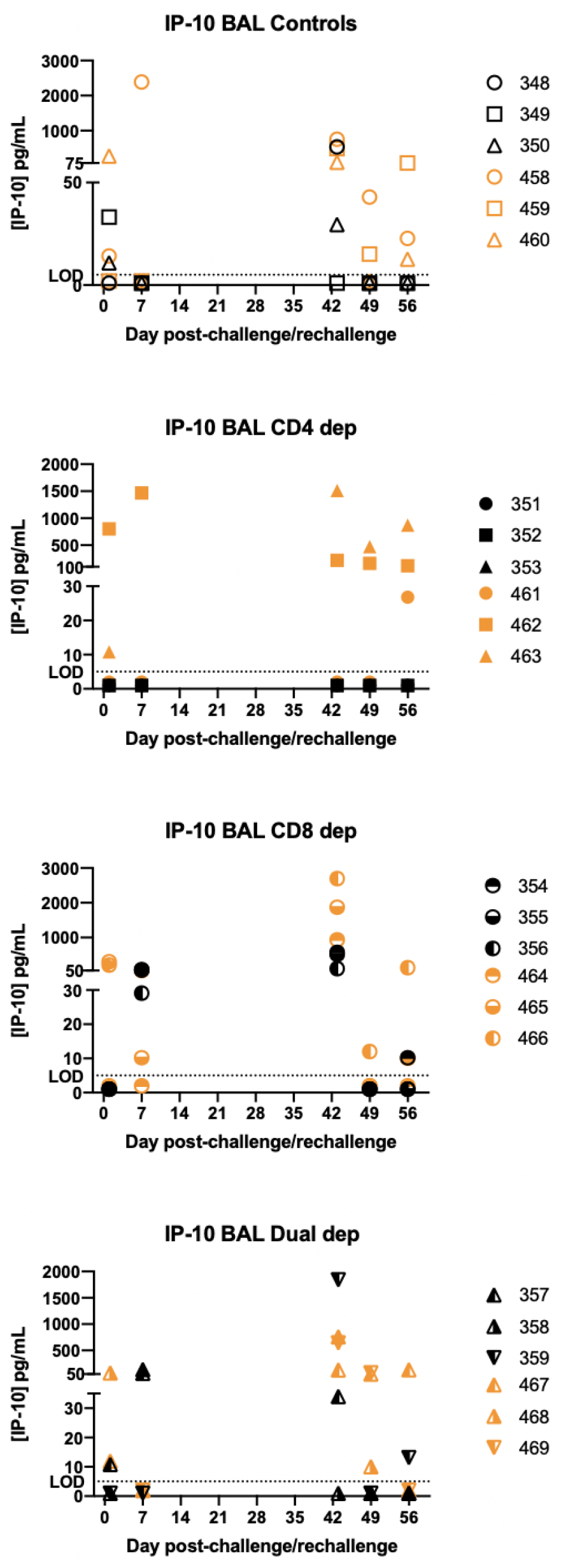
IP-10 levels in broncho-alveolar lavage fluids.

